# The pattern of Nodal morphogen signaling is shaped by co-receptor expression

**DOI:** 10.1101/2019.12.30.891101

**Authors:** Nathan D. Lord, Adam N. Carte, Philip B. Abitua, Alexander F. Schier

## Abstract

Embryos must communicate instructions to their constituent cells over long distances. These instructions are often encoded in the concentration of signals called morphogens. In the textbook view, morphogen molecules diffuse from a localized source to form a concentration gradient, and target cells adopt fates by measuring the local morphogen concentration. However, natural patterning systems often incorporate numerous co-factors and extensive signaling feedback, suggesting that embryos require additional mechanisms to generate signaling patterns. Here, we examine the mechanisms of signaling pattern formation for the mesendoderm inducer Nodal during zebrafish embryogenesis. We find that Nodal signaling activity spans a normal range in the absence of signaling feedback, suggesting that diffusion is sufficient for Nodal gradient formation. We further show that the range of endogenous Nodal ligands is set by the EGF-CFC co-receptor Oep: in the absence of Oep, Nodal ligands spread to form a nearly uniform distribution throughout the embryo. In turn, increasing Oep levels sensitizes cells to Nodal ligands. We recapitulate these experimental results with a computational model in which Oep regulates the diffusive spread of Nodal ligands by setting the rate of capture by target cells. This model predicts, and we confirm *in vivo*, the surprising observation that a failure to replenish Oep during patterning transforms the Nodal signaling gradient into a travelling wave. These results reveal that patterns of Nodal morphogen signaling are shaped by co-receptor-mediated restriction of ligand spread and cell sensitization.

## Introduction

Developing embryos often transmit instructions using morphogens, diffusible signaling molecules that induce concentration-dependent responses in target cells. In the most common conception of morphogen function, ligands spread from a localized source to form a concentration gradient^1,2^. Cells within the gradient infer their position by sensing the local ligand concentration and initiate a position-appropriate gene expression program^3,4^. Examples of gradient-driven patterning in animal embryos are plentiful; vertebrate germ layer induction^5–7^, dorsoventral organization of the neural tube^8,9^, and digit patterning^10,11^ all rely on graded profiles of signaling molecules. The biological and physical processes that set the shape of morphogen gradients are therefore of key importance to understanding developmental patterning.

Diffusion plays a central role in classical models of morphogen gradient formation. Ligand diffusion from a localized source is sufficient to create a concentration gradient that expands outward over time^12^. Adding removal of the morphogen (through degradation, internalization, or other means) to the model confers stability^13^. In such models, a steady-state gradient that does not further change in time can form^14^. The shape of this steady-state gradient reflects a balance between ligand mobility and stability. Increasing the diffusion rate lengthens the gradient, whereas faster removal shortens it^14^. Though simple, such diffusion-removal models approximate the behavior of several well-studied morphogens^15^. Recent biophysical studies have shown that fluorescently-tagged morphogens in *Drosophila*^16,17^ and zebrafish^9,18–20^ have diffusion rates consistent with known ranges of action. Similarly, receptor-mediated ligand capture provides a plausible mechanism for morphogen removal and has been shown to be a determinant of gradient range in some cases^19,21–25^.

While these simple principles seem sufficient to explain gradient formation, diffusive transport may carry inherent limitations. For example, diffusing ligands could be difficult to contain without physical boundaries between tissues^26^, and receptor saturation could preclude stable gradient formation^27^. Embryos may therefore need additional layers of control to spread signaling in a controlled fashion. Indeed, developmental signaling circuits often incorporate extensive feedback on morphogen production and sensing^3,28–30^. In these systems, signaling pattern shapes can be determined by the action of feedback rather than the biophysical properties of signaling molecules. For example, it has been argued that positive feedback on ligand production can substitute for diffusion as a mechanism of morphogen dispersal^31^. In this scheme, a cascade of short-range interactions—one tier of cells induces signal production in the next— can propagate signaling in space, even when the ligand itself is poorly diffusive. Such ‘relay’ mechanisms have been invoked to explain germ layer patterning in zebrafish^31^, as well as Wnt signal spread in micropatterned stem cell colonies^32^. Negative feedback can also shape signaling gradients, for example, by scaling patterns to fit tissue size^33,34^, restricting signaling in space^21^, or turning off pathway activity when it is no longer needed^31,35^. Due to the abundance of mechanisms that can contribute to signaling pattern shape, the mechanisms of gradient formation remain points of contention, even for well-studied morphogens.

Here, we examine the mechanism of gradient formation for the canonical morphogen Nodal. Nodals are TGFβ family ligands that function by binding to cell surface receptor complexes consisting of Type I and Type II activin receptors and EGF-CFC family co-receptors^6,7,36^. Receptor complex formation induces phosphorylation and nuclear accumulation of the transcription factor Smad2, which cooperates with nuclear cofactors to activate Nodal target genes^37^. In early vertebrate embryos, Nodal signaling orchestrates germ layer patterning: exposure to high, intermediate and low levels of Nodal correlate with selection of endodermal, mesodermal and ectodermal fates, respectively^38–41^. Nodal signaling is under both positive and negative feedback control. Nodal ligands induce the expression of *nodal* genes^42^, as well as of *leftys*^*42,43*^, diffusible inhibitors of Nodal signaling. These feedback loops are conserved throughout vertebrates and therefore appear crucial to the function of the patterning circuit^7^.

Zebrafish mesendoderm is patterned by two Nodal signals, Cyclops and Squint^7,40^. The physiologically relevant ligands are heterodimers between Cyclops or Squint and a third TGFβ family member, Vg1^44–46^. Gradient formation is initiated by secretion of Nodal ligands from the extraembryonic yolk syncytial layer (YSL), below the embryonic margin. Over time, the Nodal patterning circuit generates a gradient of signaling activity that, at the onset of gastrulation, extends approximately 6-8 cell tiers from the margin^31,47^. Mutations that markedly expand signaling range (e.g. *lefty1;lefty2*) result in profound phenotypic defects and embryonic lethality^47^. Proper development therefore relies on the generation of a correct Nodal signaling gradient.

Early studies with ectopically-expressed Nodal ligands in zebrafish supported a model of diffusive spread^48^. Direct observation of diffusion using GFP-tagged Cyclops and Squint ligands suggested short and intermediate ranges of activity, respectively^18^. In this model, the distance that ligands can diffusively travel during the ~2h prior to gastrulation is a crucial determinant of gradient range. More recently, it was argued that Nodal signal spread was driven instead by positive feedback^31^. In this model, a feedback-driven relay spreads signaling activity away from the margin, and spread is stopped by the onset of Lefty production. In contrast to the diffusion-driven model, the range of signaling is set by the properties of the feedback circuit (e.g. the time required for a cell to switch on Nodal production and delay in onset of Lefty production).

In this study, we re-examine the mechanisms that regulate Nodal signaling gradient formation in zebrafish embryos. We find that endogenous Nodal ligands can spread over a normal range in the absence of signaling feedback, suggesting that diffusion is sufficient for gradient formation. Unexpectedly, we discover that the EGF-CFC co-receptor Oep is a potent regulator of the range of both Cyclops and Squint; in mutants lacking *oep*, Nodal ligands achieve a near-uniform distribution throughout the embryo. We also find that Oep, though traditionally regarded as a permissive signaling factor, sets cell sensitivity to Nodal ligands. We incorporate these observations into a mathematical model for Nodal signal spread and predict that replenishment of Oep by zygotic expression is required for gradient stability. Finally, we verify a surprising prediction of the model: in zygotic *oep* mutants, which cannot replace Oep after it has been degraded, Nodal signaling propagates outward from the margin as a traveling wave. These findings illustrate how the embryo uses an unappreciated property of Oep—regulation of the rate of ligand capture— to set the range and intensity of the Nodal signaling gradient.

## Results

### The Nodal signaling gradient forms in the absence of feedback

The Nodal signaling gradient may reflect the diffusive properties of Nodal ligands secreted from the YSL or the action of signaling feedback. To characterize the contribution of diffusion specifically, we set out to visualize the Nodal gradient in mutants that lack signaling feedback altogether. This goal presented two key challenges. First, endogenous Nodal ligands have not been successfully visualized by antibody staining or fluorescent tagging in zebrafish. Second, knocking out the full complement of all known Nodal feedback regulators— e.g. *lefty1, lefty2, cyclops, squint, dpr2*^*49*^, etc.— in combination is impractical. To address these two limitations, we developed a ‘sensor’ cell assay (Fig. 1A). In this approach, we transplant Nodal-sensitive (‘sensor’) cells from a *gfp*-injected donor embryo to the margin of a host that is Nodal-insensitive and therefore lacks feedback. We then visualize signaling in the sensor cells by immunostaining for phosphorylated Smad2 (pSmad2) and GFP. Because host cells cannot respond to Nodal, they cannot modulate signal spread by either positive or negative feedback. The sensor cells ‘report’ on their local Nodal concentration via pSmad2 staining intensity, enabling us to sample the activity of endogenous, untagged ligands. For the experiments described here, we use sensor cells from M*vg1* donors. These cells are Nodal sensitive but cannot produce functional Nodal ligands and therefore cannot spread signaling via positive feedback^44^.

**Fig. 1.**
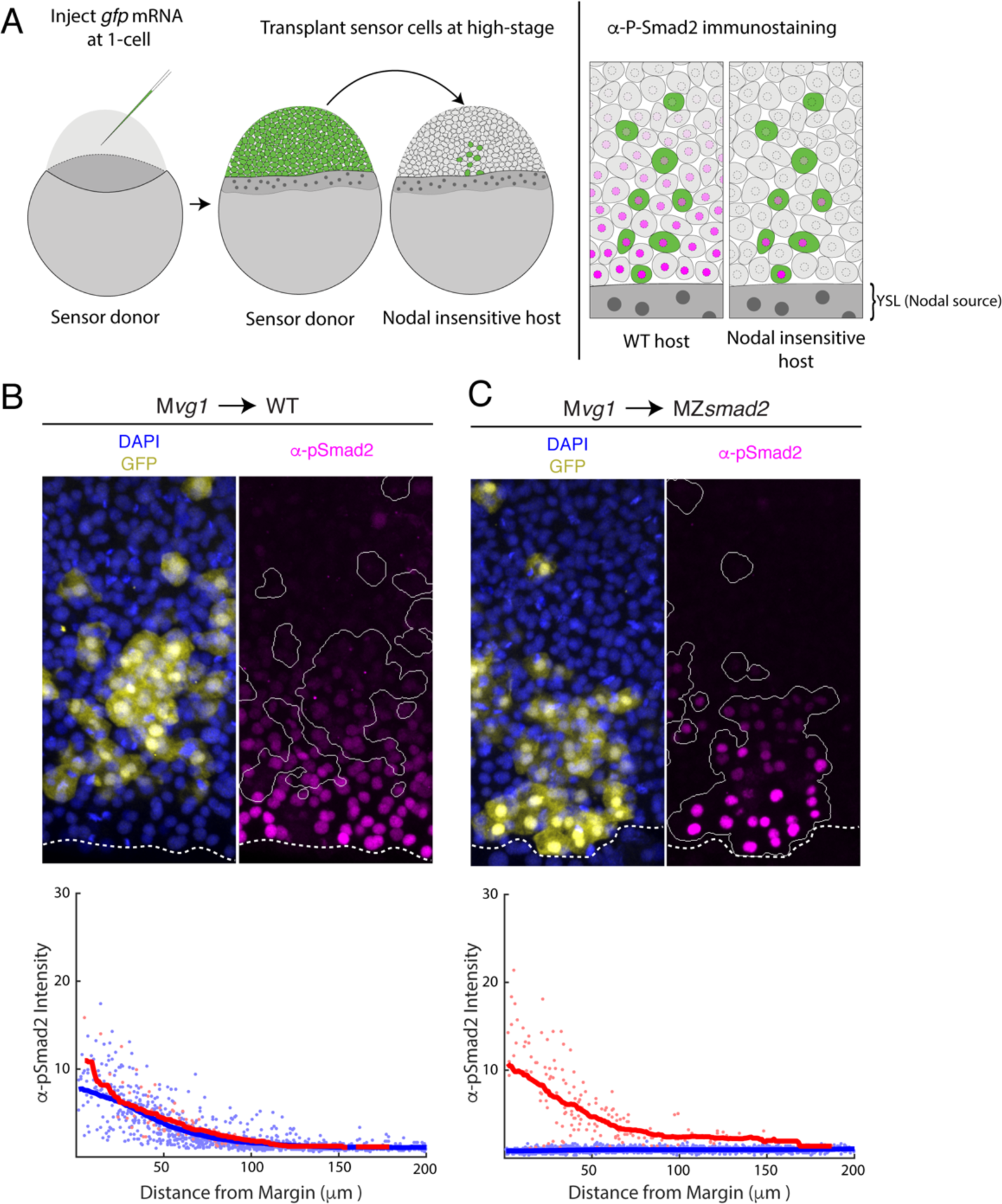
Nodal gradient formation in the absence of feedback. **A)** Schematic of sensor cell assay. M*vg1* donor embryos were marked by injecting *gfp* mRNA at the 1-cell stage. At high stage, just before the onset of Nodal signaling, GFP-marked sensor cells were transplanted from the animal pole of the donor to the margin of a Nodal-insensitive host. At 50% epiboly, embryos were fixed and immunostained for GFP and Nodal signaling activity (α-pSmad2). Imaging of chimeric embryos (far right) enables inference of the gradient shape from α-pSmad2 staining (magenta) in sensor cells (green). Because host embryos lack the ability to respond to Nodal, YSL-derived Nodal ligands are responsible for the shape of the Nodal signaling gradient. **B)** Control visualization of the Nodal signaling gradient in wild-type hosts using a sensor cell assay. Upper panel; M*vg1* sensor cells (yellow) were transplanted to the margin of a wild-type host. Nodal signaling was visualized by α-pSmad2 staining (magenta), and sensor cell boundaries were segmented with an automated pipeline (white curves). YSL boundaries are marked with dashed white curves. Lower panel; quantification of staining intensity in host (blue) and sensor (red) cells across replicate embryos. Nuclei were segmented from DAPI signal using an automated analysis pipeline implemented in MATLAB. Sensor and host cells were identified as being clearly GFP positive or negative, respectively. Solid curves represent sliding window averages. **C)** Sensor cell assay in MZ*smad2* host embryos. Upper panel; GFP-marked M*vg1* sensor cells (yellow) were transplanted to the margin of MZ*smad2* host embryos. Nodal signaling was visualized with α-pSmad2 staining (magenta). Sensor cell boundaries are marked with white outlines, and YSL boundaries are marked with dashed white curves. Lower panel; quantification of host (blue) and sensor (red) cell staining intensities were carried out as in (B).

We first applied this approach to MZ*smad2* host embryos, which lack all Nodal signaling; Smad2 is required to activate Nodal-dependent gene expression, and zebrafish MZ*smad2* embryos phenocopy mutants lacking Nodal ligands^50^. We verified MZ*smad2* embryos lack pSmad2 (Fig. S1) but continue to express *cyclops* and *squint* in the YSL (Fig. S2). Expression of both Nodals was excluded from the blastoderm, confirming that these mutants are incapable of Nodal autoregulation (Fig. S2). M*vg1* sensor cells transplanted into MZ*smad2* mutants exhibit clear Nodal signaling activity several cell tiers from the margin (Fig. 1C, upper panel), while signaling was completely absent in host cells. Control transplants into wild-type embryos confirmed that the sensors accurately reported on their signaling environment (Fig. 1B, upper panel); Mvg1 sensors exhibited α-pSmad2 staining intensity similar to their wild-type neighbors, and quantification of staining across replicate embryos revealed similar signaling gradients for host and sensor cells (Fig. 1B, lower panel; blue and red points, respectively). Quantification of staining in MZ*smad2* hosts (Fig. 1C, lower panel) revealed a Nodal signaling gradient similar in range to that of wild-type controls (half-distances of 45 and 37 μm for MZ*smad2* and wild type, respectively). Together, these experiments suggest that YSL-derived Nodal ligands can form a gradient of normal range without help from signaling feedback.

### Nodal signaling range is expanded in the absence of Oep

The above results support a model in which diffusion drives Nodal spread. However, it remains unclear how the embryo sets the range of ligand dispersal. Biophysical studies with GFP-tagged Nodals suggest that ligand mobility may be hindered by interaction with extracellular factors. Measured diffusion rates for both Cyclops and Squint are >10-fold lower than for free GFP^18^; however, no factors that explain hindered mobility of endogenous ligands have been identified. Cell surface receptor complexes are clear candidates for this role^51^. Transient ligand capture or receptor-mediated endocytosis could constrain the gradient^14^, and receptors have been shown to regulate gradient range for other signals^21,22,25,52^.

To test whether receptor complex components regulate the range of Nodal signaling, we performed sensor cell transplants in embryos lacking the essential Nodal co-receptor Oep (MZ*oep* mutants^36^). We found that M*vg1* sensor cells detected Nodal activity over a dramatically longer range in MZ*oep* hosts than in wild-type controls (*cf.* Figs. 2A,B). Indeed, transplanting sensor cells to the animal pole revealed that Nodal ligands can be detected throughout the embryo when Oep is absent (Figs. 2D,E). To test whether loss of Oep affects both Nodal ligands similarly, we performed sensor cell assays in MZ*oep*;*sqt* and MZ*oep*;*cyc* double mutants. Loss of Oep led to an expanded range of action for both Cyclops (i.e. in MZ*oep;sqt* mutants) and Squint (i.e. in MZ*oep*;*cyc* mutants), and the signaling ranges in both double mutants were comparable to what we observe in the MZ*oep* single mutant (Fig. S3).

**Fig. 2.**
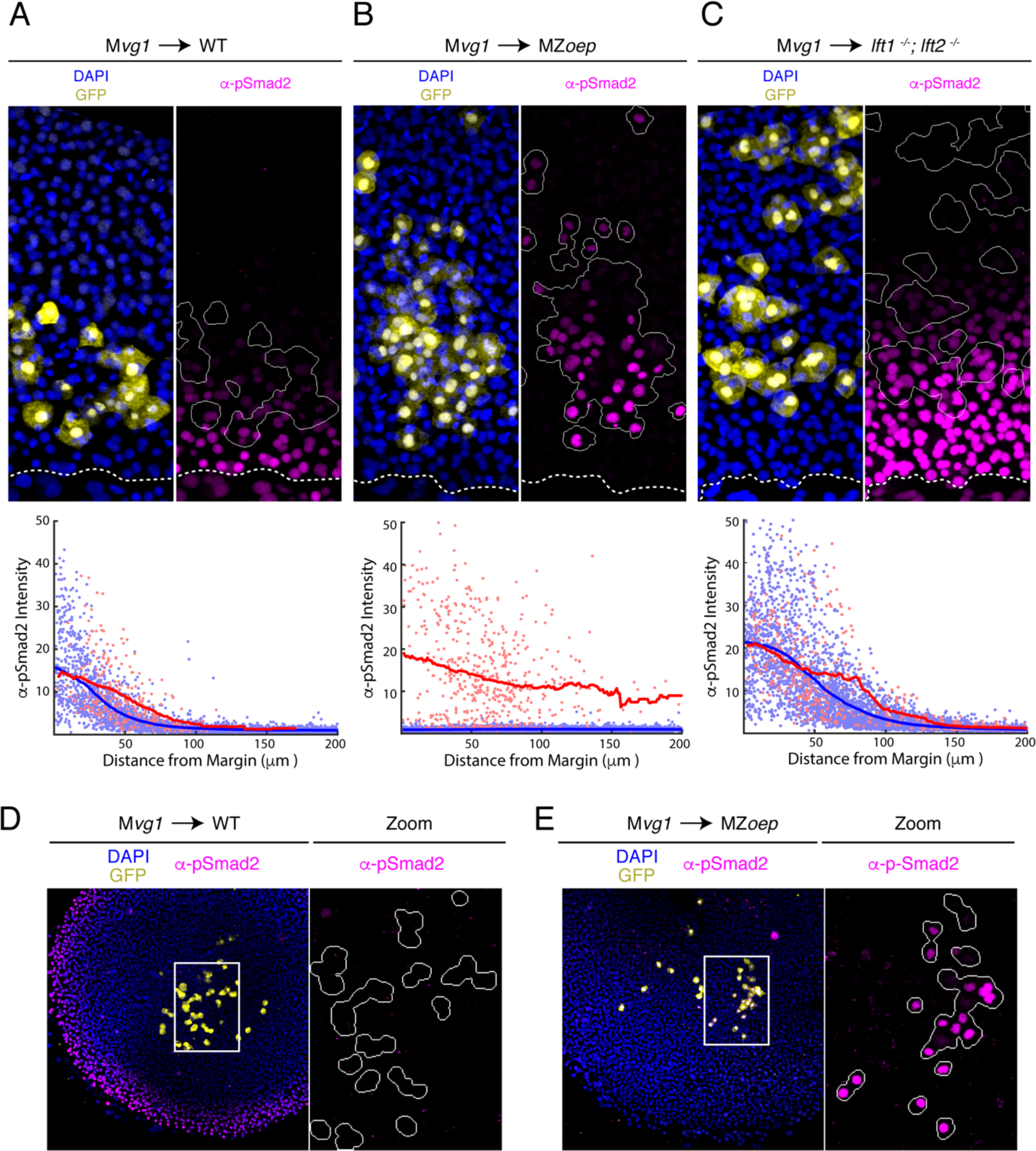
The Nodal gradient is expanded in MZ*oep* mutants. **A-C)** Sensor cell assay and gradient quantifications in (A) wild type, (B) MZ*oep* and (C) *lft1*^*−/−*^;*lft2*^*−/−*^ embryos. M*vg1* sensor cells were marked with GFP (yellow) and transplanted to the margin of host embryos. Nodal signaling activity is measured by α-pSmad2 immunostaining (magenta). YSL boundaries are marked with dashed curves and sensor cell boundaries are outlined in solid white in all α-pSmad2 panels. Gradient quantifications for each experiment are below images; host and sensor cell staining intensities are plotted as blue and red points, respectively. Sliding window averages are plotted as solid curves. **D)** Left panel; M*vg1* sensor cells (yellow) were transplanted directly to the animal pole of a wild-type host. The endogenous Nodal signaling gradient is visible at the embryonic margin (magenta). White box highlights region expanded for detail view in right panel. Right panel; Nodal signaling activity is absent in both host and sensor cells. **E)** Left panel; M*vg1* sensor cells (yellow) were transplanted to the animal pole of an MZ*oep* embryo. Nodal signaling is absent at the embryonic margin. White box highlights region expanded in the right panel. Right; sensor cells detect Nodal at the animal pole (magenta).

The magnitude of gradient expansion in MZ*oep* mutants is remarkable when compared with the effect of other mutations that alter Nodal signaling range. For example, the signaling gradient is expanded in *lefty1;lefty2* mutant embryos, which lack negative feedback on Nodal signaling^47^ (Fig. 2C). This degree of gradient expansion causes profound phenotypic defects in these mutants but is mild compared to our observations in MZ*oep* embryos (*cf*. Figs. 2B,C). This difference is particularly notable given that a positive feedback relay may propagate signaling in *lefty* mutants. In MZ*oep* mutants, which lack positive feedback on signaling, the gradient expansion reflects only changes to the range of YSL-derived Nodal ligands. Together, these results demonstrate that receptor complexes constrain the spread of Nodal signals.

### Oep regulates the range and intensity of Nodal signaling through ligand capture

EGF-CFC proteins such as Oep are typically regarded as permissive factors for Nodal signaling. Oep facilitates the assembly of receptor-ligand complexes but is not thought to regulate signaling beyond conferring competence^53^. However, our finding that Nodal ligand range is expanded in the absence of Oep suggests that it has unappreciated regulatory roles. The simplest way to accommodate this result is to stipulate that Oep levels set the rate of capture of diffusing Nodal ligands. Through this mechanism, Oep could control the range of Nodal activity by regulating the rate of receptor-mediated endocytosis (i.e. the effective ligand degradation rate). This model makes two testable predictions. First, increasing Oep levels should enhance cell sensitivity to Nodal ligands by facilitating capture by receptor complexes. Second, increasing Oep levels should reduce the range of Nodal signaling by increasing the effective degradation rate.

To test whether Oep regulates cell sensitivity, we asked whether overexpressing *oep* in sensor cells increases their responsiveness to endogenous Nodals. We transplanted cells from M*vg1* embryos injected with *oep* and *gfp* mRNAs or with *gfp* alone to the margin of wild-type embryos and immunostained for GFP and pSmad2. Sensors with increased Oep levels stained more brightly for pSmad2 than neighboring host cells (Fig. 3B), while sensors injected with *gfp* alone matched the behavior of their neighbors (Fig. 3A). Interestingly, we found that the +*oep* sensors detected Nodal further from the margin than the host cells, suggesting that the Nodal ligand gradient extends beyond the domain of detectable signaling in normal embryos (Fig. 3B). We note that the increased sensitivity of the +*oep* sensors does not reflect the action of hyperactive positive feedback on Nodal production, as M*vg1* cells are incapable of producing functional Nodal ligands. These results suggest that, in addition to being required for signaling competence, Oep regulates sensitivity to Nodal ligands.

**Fig. 3.**
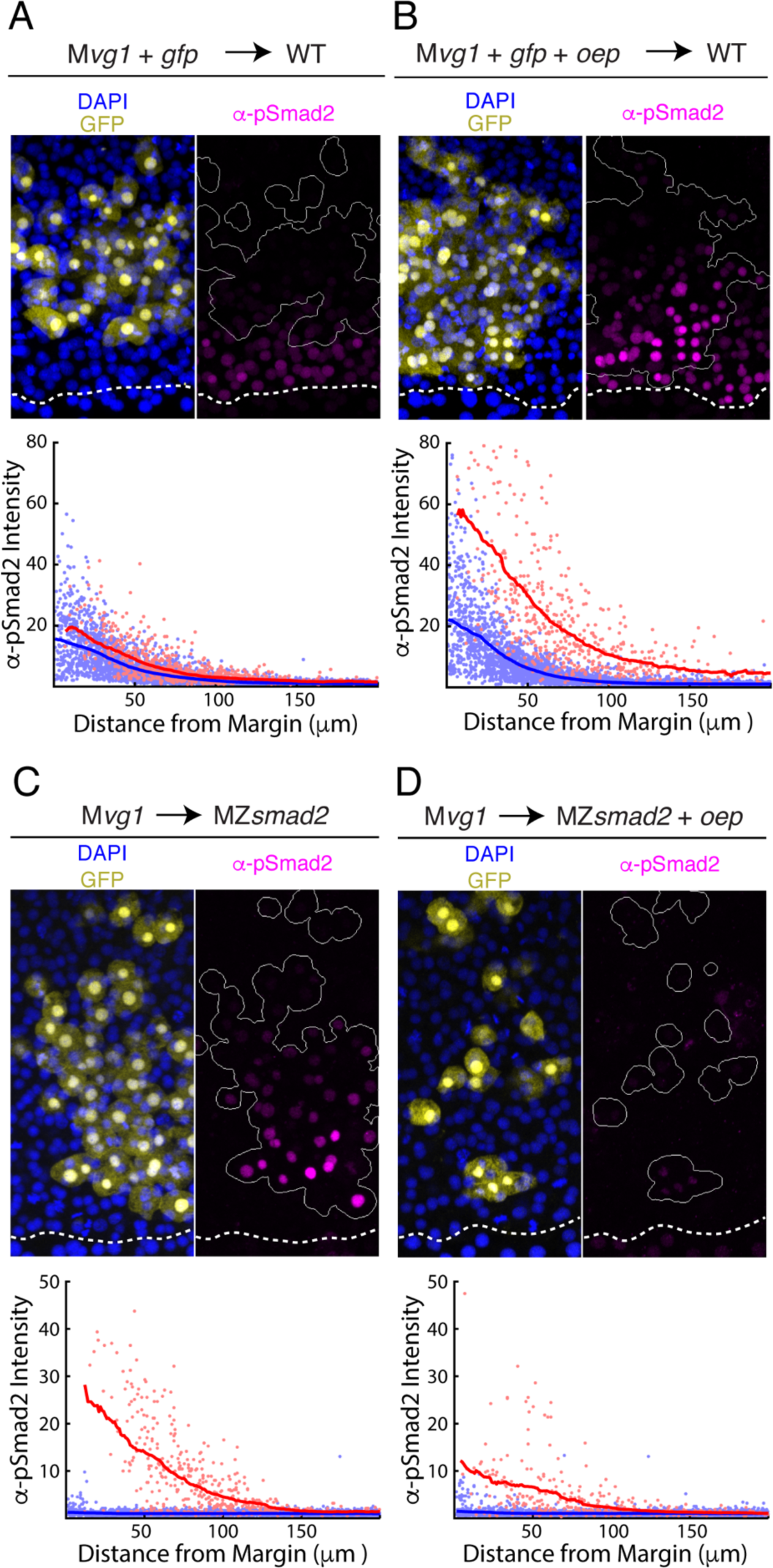
Oep levels regulate Nodal ligand capture. **A-B)** Oep overexpression increases sensitivity to Nodal ligands. **A)** Upper panel: control transplant of GFP-marked M*vg1* sensor cells (yellow) to the margin of wild-type hosts. Nodal signaling activity was measured by α-pSmad2 immunostaining (magenta). In all panels, YSL boundaries are marked with dashed white curves, and sensor cells have been outlined in solid white in all α-pSmad2 panels. Lower panel: quantification of Nodal signaling in sensor (red) and host cells (blue) across replicate embryos. Sliding window averages are plotted as solid curves. **B)** Upper panel: transplant of sensor cells from an M*vg1* donor injected with *gfp* and 110 pg *oe*p mRNA at the 1-cell stage to the margin of wild-type hosts. Sensor cells (yellow) exhibit enhanced Nodal signaling activity (magenta) compared to their host-derived neighbors. Lower panel; staining of host (blue) and sensor (red) cells was quantified as in (A). **C-D)** Oep overexpression restricts Nodal spread. **C)** Upper panel: sensor cell measurement of the Nodal gradient in MZ*smad2* embryos. M*vg1* sensor cells were marked with GFP (yellow), and Nodal signaling activity was measured by α-pSmad2 immunostaining (magenta). Lower panel: quantification of Nodal signaling in sensor (red) and host cells (blue) was quantified as in (A). **D)** Upper panel: M*vg1* sensor cell measurement of the Nodal signaling gradient in MZ*smad2* hosts injected with 110 pg *oep* mRNA at the 1-cell stage. Lower panel; gradients were quantified as in (A).

To test whether Oep levels regulate Nodal range, we asked whether overexpression of *oep* could restrict signaling. We performed sensor cell assays in MZ*smad2* hosts injected with *oep* mRNA at the 1-cell stage. Overexpression of oep indeed reduced the range and intensity of Nodal signaling (Fig. 3D) when compared with uninjected hosts (Fig. 3C). We note that the choice of MZ*smad2* hosts was important for interpretation of the experiment. As Oep sensitizes cells to Nodal ligands, increasing expression in signaling-competent host embryos could lead to increased signaling by triggering Nodal positive feedback. Nodal signaling is disabled downstream of the receptor in MZ*smad2* mutants, allowing us to specifically test Oep’s role in regulating ligand range without this confound. Together, these results suggest that, by facilitating capture of Nodal ligands, Oep regulates range and intensity of the Nodal activity gradient.

### A simple model incorporating Oep-Nodal interaction reproduces experimental observations

We formulated a simple mathematical model of Nodal gradient formation to explore whether Oep-mediated capture of diffusing Nodal ligands is sufficient to explain our experimental data (Fig. 4A). In the model, Nodal is secreted at a constant rate at one end of a 2-dimensional tissue and diffuses freely until it is captured by a free receptor complex. We stipulate that ligand-receptor association follows pseudo first-order kinetics (i.e. that the free receptor concentration can be regarded as constant) and that internalization of receptor-ligand complexes is also first-order. To track integration of signaling activity, we also incorporate phosphorylation of Smad2 with a rate proportional to ligand-receptor complex concentration. Where possible, parameter values were taken from the literature. A summary of the rates used in simulations is presented in Table 1.

**Fig. 4.**
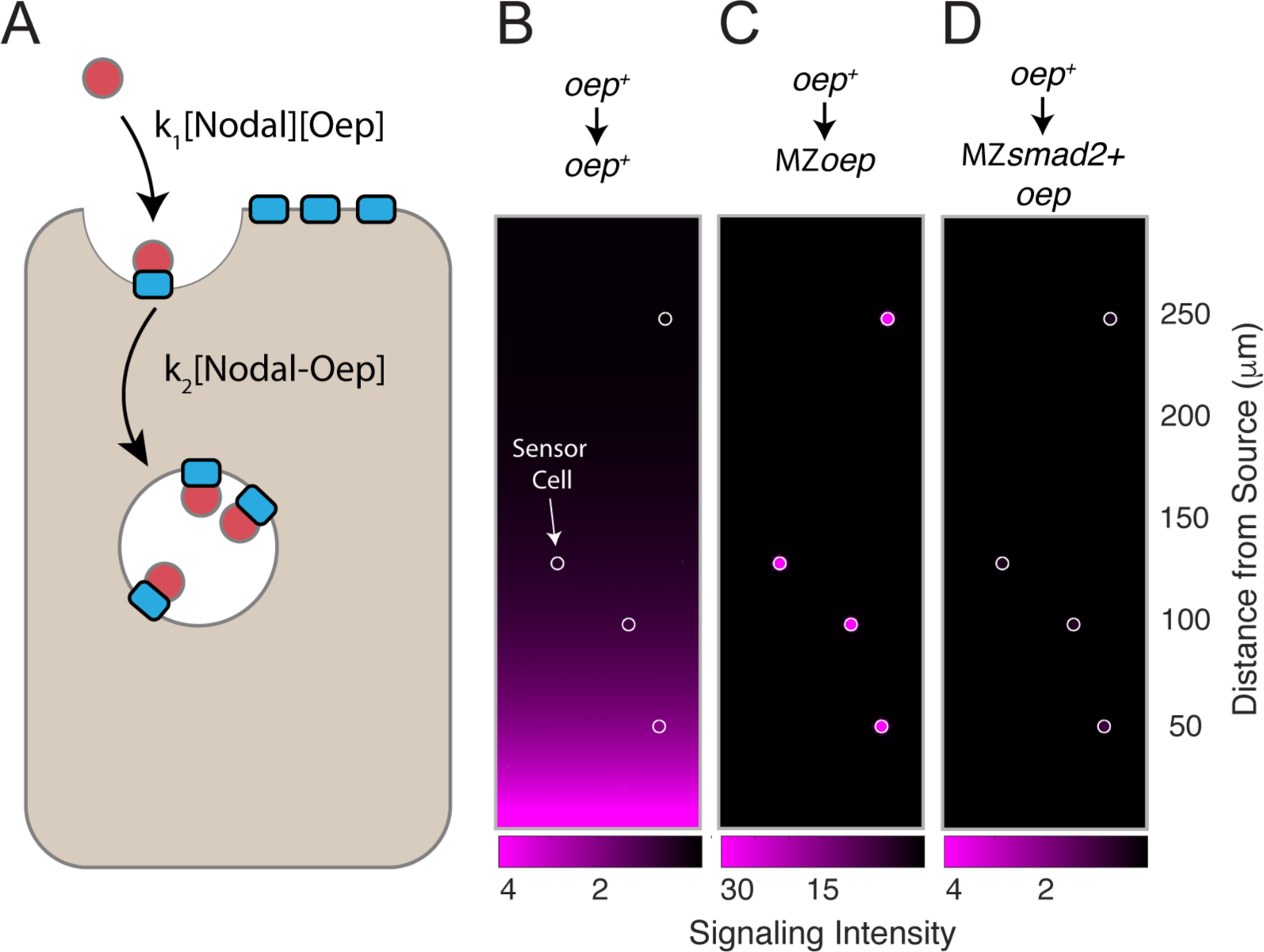
A simple model of Nodal diffusion and capture reproduces experimental observations. **A)** Schematic of Nodal diffusion-capture model. Simulations were performed on a two-dimensional tissue of 100 μm x 300 μm. Nodal molecules are secreted at a constant rate from a localized source at one boundary of the tissue (i.e. 0 < x < 5 μm) and diffuse freely until capture by cell surface receptors (‘Oep’). Ligand-receptor complexes are removed from the system by internalization. To track signaling activity, Smad2 phosphorylation is simulated with rate proportional to the concentration of receptor-ligand complexes. **B-D)** Simulation of transplant experiments. In each simulation, the behavior of sensor cells (white outlines) is compared with the behavior of the host embryo (remainder of tissue). Parameters were independently set for host and sensor regions, allowing for simulation of experiments with mutations and overexpression. Signaling activity (i.e. [pSmad2]) is plotted in magenta. **B)** Wild-type gradient simulation. Sensor cells with normal Oep levels are transplanted into a host with normal Oep levels. A stable gradient forms, and signaling is identical in sensor cells and neighboring regions. **C)** Gradient expansion in MZ*oep* mutants. Sensor cells contain normal Oep levels, but host cells lack Oep. Sensor cells detect ligand throughout the tissue. **D)** Gradient contraction with *oep* overexpression. Sensor cells contain normal Oep levels, whereas host cells lack Smad2, but overexpress *oep*. Signaling is absent in the host tissue—due to lack of Smad2— but elevated receptor expression restricts Nodal spread to the sensors.

This simple model reproduces a signaling gradient with a scale and shape consistent with our observations in wild-type embryos (Fig. 4B). To reproduce our experimental data, we simulated sensor cell assays (Figs. 4B-D, sensor cells highlighted with white outlines). Expansion of the Nodal ligand gradient in MZ*oep* mutants can be reproduced by simulating ‘hosts’ with the receptor concentration set to zero (Fig. 4C). Similarly, restriction of signaling range via *oep* overexpression could be captured by increasing receptor levels in host cells, but not in the sensors (Fig. 4D). A model in which Nodal capture rate is set by Oep concentration can therefore reproduce our major experimental findings.

### Loss of Oep replenishment transforms Nodal signaling dynamics

The simplified model presented above assumes that free receptor cannot be depleted by ligand binding. While convenient, this condition may be difficult for the embryo to achieve in practice. For example, maintaining receptors at high concentration would preclude depletion but could also prevent ligand from traveling long distances before capture. Another way for the embryo to avoid depletion would be to continually replace receptor components as they are consumed by ligand binding. To explore the role of receptor complex replacement in gradient formation, we explicitly incorporated receptor production and degradation into the model (Fig. 5A).

**Fig. 5.**
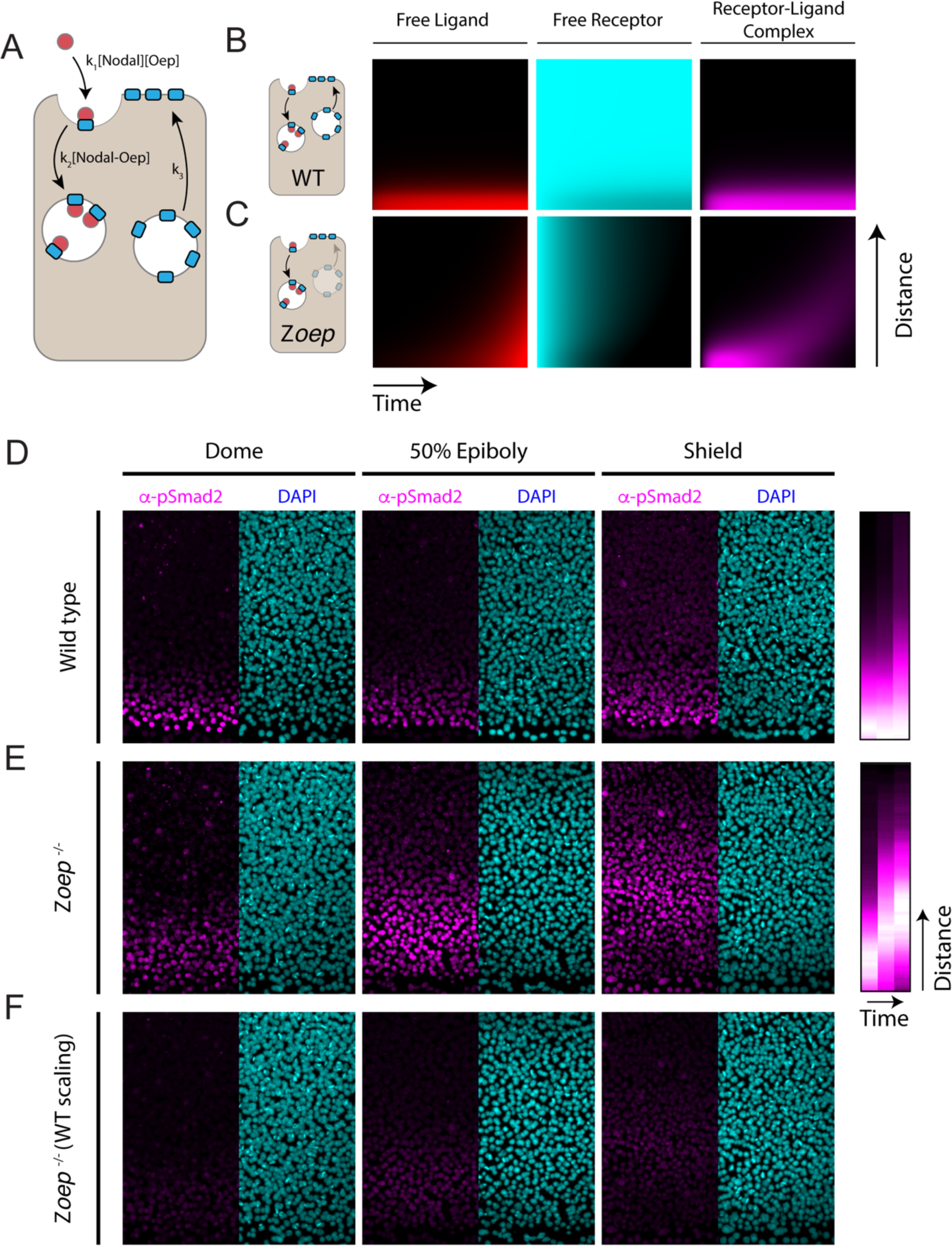
Loss of Oep replacement destabilizes the Nodal signaling gradient. **A)** Schematic of model incorporating production and consumption of receptors. Simulations presented here were performed on a one-dimensional tissue with length 300 μm. Oep replacement is assumed to be constant with rate k_3_, and Oep removal reflects a combination of constitutive and ligand-dependent endocytosis. In panels A and B, simulations are presented as kymographs; each image column shows the state of the system with the source at the bottom and animal pole at the top. Time proceeds from left to right. **B)** Simulation of a wild-type gradient. With continual receptor replacement, the system achieves an exponential steady state gradient with length scale set by the ligand diffusion rate and receptor abundance. The free ligand, free receptor and receptor-ligand complex concentrations are plotted from left to right in red, cyan and magenta, respectively. **C)** Simulation of gradient formation in a zygotic *oep* mutant. Simulation details are identical to (B), but with receptor replacement rate (k_3_) set to zero. The system fails to establish a steady state due to gradual consumption and degradation of receptors. Over time, the Nodal ligand gradient expands (red) to drive a propagating wave of signaling activity (i.e. receptor occupancy, magenta). **D)** Time course of Nodal signaling activity in wild-type embryos. Representative α-pSmad2 (magenta) and DAPI (cyan) are shown for dome, 50% epiboly and shield stages (left, middle and right panels, respectively). Quantification of signaling gradients across replicates (far right) shows the establishment of the signaling gradient. **E)** Time course of Nodal signaling activity in zygotic *oep* mutants. Over time, the signaling pattern evolves from a gradient (dome stage) to a band displaced far from the margin (shield) as the wave travels outward. Quantification of signaling gradients across replicates (far right) illustrates the outward propagation of signaling. **F)** Time course of Nodal signaling activity in zygotic *oep* mutants presented with pixel scaling equal to that used in (D). In accord with simulations, the wave of signaling propagates with a lower intensity than signaling at the margin of WT embryos.

Simulations incorporating receptor production and consumption generate stable exponential gradients (Fig. 5B) with length scales comparable to our measurements in zebrafish embryos. To test the consequences of abolishing co-receptor replacement, we simulated gradient formation in a system that begins with a finite supply of free receptors that are not replaced. This change results in a surprising transformation of Nodal signaling dynamics; simulations with finite co-receptor supply generate a traveling wave of Nodal signaling that propagates outward from the ligand source (Fig. 5C, magenta). These dynamics reflect the gradual consumption of co-receptors due to ligand binding and subsequent endocytosis (Fig. 5C, cyan). Initially, when co-receptor is plentiful, the source generates a decaying gradient of signaling. Over time, receptors close to the source are depleted, allowing Nodal ligands to rapidly traverse this space, ultimately reaching a new population of sensitive cells. These simulations raise the possibility that co-receptor replenishment is a key determinant of the Nodal gradient shape.

To test this possibility, we measured Nodal signaling patterns in zygotic *oep* mutants (Z*oep*). This background reproduces the key assumptions of the model above: Z*oep* mutants begin with a finite supply of maternally-provided *oep* mRNA but cannot express additional *oep* from the zygotic genome. We performed α-pSmad2 immunostaining in wild-type and Z*oep* mutant embryos at three timepoints following the initiation of Nodal secretion (dome, 50% epiboly and shield stages). Consistent with previous observations, the wild-type Nodal signaling profile monotonically decreases from the margin, decaying to background over ~8 cell tiers (Fig. 5D). Strikingly, in Z*oep* mutants, Nodal signaling is restricted to the margin at dome stage (Fig. 5D, left), but propagates outward to form a broad band of signaling by shield stage (Fig. 5D, right). As predicted by the model, loss of co-receptor replacement by zygotic expression thus transforms a steady-state exponential gradient into a wave of Nodal signaling that propagates toward the animal pole. We note that, in accordance with model simulations, overall signaling intensity is lower in Z*oep* mutants due to lower overall co-receptor levels (Fig. 5F). These results highlight the importance of continued co-receptor replacement in shaping the pattern of Nodal signaling.

## Discussion

In this study, we set out to identify mechanisms that determine the Nodal signaling gradient range. We find that endogenous Nodals secreted from the YSL can drive signaling over a normal range in the absence of feedback (Fig. 1). We go on to demonstrate that expression of Oep, a Nodal co-receptor, regulates the spread (Fig. 2) and potency (Fig. 3) of Nodal ligands. We propose a computational model that explains the Nodal signaling gradient in terms of free ligand diffusion and binding to cell surface receptor complexes (Fig. 4). In this description, Oep regulates the range of ligand spread and sensitivity of embryonic cells by setting the rate of ligand capture. This simple model accommodates our main observations—gradient formation without feedback, increased signaling range in co-receptor mutants, and restricted range with increased co-receptor expression— and predicts the surprising Nodal signaling wave in zygotic *oep* mutants (Fig. 5).

Diffusion has long been regarded as an attractive mechanism for signal dispersal in tissues^13^. Indeed, signaling patterns consistent with simple diffusion-degradation mechanisms—e.g. single-exponential gradients with length scales of ~10-100 um— are common in developing tissues^15^. Viewed in this light, the regulatory complexity of developmental patterning circuits is striking; if diffusion is sufficient to generate observed signaling patterns, why are a plethora of co-factors and extensive feedback loops so common? One possible answer is that diffusion carries inherent disadvantages. For example, it has been argued that diffusible ligands would be impractical to contain without physical boundaries^26^, and that diffusion-driven gradients would not be a reliable source of positional information^54^. We and others have proposed feedback-centered Nodal patterning models that offer a way around these dilemmas^18,31,55^. However, it has not been possible to clearly test whether feedback is required for the dispersal of endogenous ligands. This study is the first to examine the shape of the Nodal signaling gradient in the absence of feedback. To our surprise, we found that a gradient of approximately normal range and shape can form even when feedback is disabled. Indeed, it seems that our observations can be largely explained by a model that relies on diffusion and capture to form the Nodal gradient.

Our study also identifies new roles for EGF-CFC co-receptors in Nodal patterning. Oep has been traditionally regarded as a permissive factor for signaling^53^; it facilitates Nodal association with Activin receptors^56–58^, but was not thought to regulate gradient shape or cell sensitivity^36,53^. Our observations suggest that— similar to receptors for Dpp^22^, Hh^21^ and Wg^25^— Oep is a key determinant of the mobility and potency of its cognate ligand. Indeed, far from being a bystander in gradient formation, Oep is one of the strongest regulators of Nodal range yet discovered. This finding also suggests a potential explanation for a key feature of the Nodal patterning circuit: differential diffusivity between Nodal ligands and Lefty proteins. GFP-tagged Cyclops and Squint diffuse substantially slower than free GFP, whereas tagged Lefty proteins diffuse rapidly^18^. This feature of Nodal ligands is consistent with a hindered diffusion model in which interactions with immobile binding partners leads to a slow ‘effective’ diffusion rate, even if free molecules diffuse rapidly^18^. Our data raise the possibility that the differential diffusivity of Nodal and Lefty proteins originates in rates of capture by available receptor complexes.

Oep-mediated ligand capture and signaling sensitization results in short-range enhancement and long-range inhibition of Nodal signaling: close to the Nodal source, Oep binds Nodal and stimulates signaling, whereas far from the source, little Nodal is available due to Oep-mediated capture close to the source. The Nodal signaling factor Oep thus has a function reminiscent of the Nodal inhibitor Lefty. Lefty is produced at the margin, but diffuses rapidly to inhibit Nodal signaling far from the source. A common theme for Nodal regulators is therefore to restrict Nodal signaling to a domain near the ligand source. This theme may reflect diffusion’s role in the propagation of Nodal signals; in the absence of physical boundaries, the Nodal patterning circuit may have to provide mechanisms for ligand containment to the intended tissues.

Our results suggest that the embryo’s strategy for replenishing Oep is a key point of control over the signaling pattern. We found that, without this replacement, the Nodal signaling pattern is qualitatively transformed from a stable gradient into a propagating wave. Interestingly, a signaling wave of this type was predicted in a theoretical study of morphogen gradient formation by Kerzsberg and Wolpert^27^. In fact, they used this phenomenon to argue that receptor saturation would make stable gradients difficult to achieve by diffusive transport. Our results suggest that consumption of receptors can create precisely this type of unstable behavior, but that the embryo achieves a stable gradient through continual turnover of the receptor pool. Though not employed during mesendodermal patterning, this phenomenon could provide a simple means of repurposing the Nodal patterning circuit to create dynamic waves of signaling in other contexts. We speculate that the precise dynamics of Oep replacement could contribute interesting functions to the patterning system. For example, signaling-dependent receptor expression could confer robustness to fluctuations in source-derived morphogen production^59,60^.

The surprising dispensability of positive feedback for gradient formation parallels our recent findings on the role of negative feedback in Nodal patterning^47^. In that work, we showed that Lefty-mediated feedback—despite its extensive conservation across animals— was dispensable for normal development in zebrafish. Lefty was instead required for robustness; intact feedback loops enabled the embryo to correct exogenous perturbations to signaling. This raises the intriguing possibility that Nodal positive feedback serves a similar purpose. Though dispensable for gradient formation *per se*, positive feedback may help to ensure that a gradient of the appropriate shape and intensity forms even in the face of mutations, environmental insults or signaling noise.

## Supporting information

Supplemental Table (smFISH Probe Sequences)

## Acknowledgments

This research was supported by the National Institutes of Health (R37GM056211 to AFS, K99-HD097297-01 to NDL, T32GM080177 training grant supported ANC), the Arnold and Mabel Beckman Foundation (postdoctoral fellowship to NDL), the NSF (GRFP DGE1745303 to ANC), a Simmons Family Imaging Award (to ANC), and the Damon Runyon Cancer Research Foundation (postdoctoral fellowship to PBA). NDL, ANC and AFS conceived the project and designed experiments; NDL and ANC performed the experimental work and analysis. NDL and ANC wrote the paper, with input from AFS. PBA identified the pSmad2 antibody used for immunostaining. We thank Kaitlyn Webster and Kellee Siegfried for the generous gift of *dmrt1* mutant zebrafish. We thank Doug Richardson and the Harvard Center for Biological Imaging for microscopy infrastructure and support. We thank Jeffrey Farrell, Katherine Rogers, Harold McNamara, and P.C. Dave P. Dingal for helpful comments on the manuscript.

## Materials and Methods

### Genotyping

Genomic DNA was isolated via the HOTSHOT method from either excised adult caudal fin tissue or individual fixed embryos^61^. Genotyping was carried out via PCR under standard conditions followed by restriction enzyme digest when appropriate. For brevity, allele designations were omitted in the rest of the text.

#### lefty1^a145^

The *lefty1*^*a145*^ allele contains a 13-base-pair deletion that destroys a PshAI restriction site and was detected as in^47^.

#### lefty2^a146^

The *lefty2*^*a146*^ allele contains an 11-base-pair deletion and was detected as described^47^.

#### squint^cz35^

The *squint*^*cz35*^ allele contains a ~1.9 kb insertion and was detected as in (Feldman et al 1998).

#### cyclops^m294^

The *cyclops*^*m294*^ allele contains a single nucleotide polymorphism (SNP) that destroys an AgeI restriction site and was detected as described^62^.

#### oep^tz57^

The *oep*^*tz57*^ allele contains a SNP that introduces a Tsp45I restriction site^53,63^. The allele was detected via PCR amplification with primers *AC102* and *AC103* flanking the SNP followed by Tsp45I digestion overnight. A wild-type allele yields an undigested band of 285 bp, while a mutant allele yields bands of 140 bp and 145 bp.

#### vg1^a165^

The *vg1*^*a165*^ allele contains a 29 bp deletion and was detected as described^44^.

#### smad2^vu99^

The *smad2*^*vu99*^ allele contains a SNP that introduces a BtsCI restriction site^50^. The allele was detected via PCR amplification with primers *NL-89* and *NL-91* flanking the SNP followed by BtsCI digestion overnight. A wild-type allele yields an undigested band of 298 bp, while a mutant allele yields bands of 221 bp and 77 bp.

### Zebrafish husbandry and fish lines

Fish were maintained per standard laboratory procedures^64^. Embryos were raised at 28.5ºC in embryo medium (250 mg/L Instant Ocean salt, 1 mg/L methylene blue in reverse osmosis water adjusted to pH 7 with NaHCO_3_) and staged according to a standard staging series^65^. Wild-type fish and embryos represent the TLAB strain. *Lefty1, lefty2, squint, cyclops, oep*, and *vg1* mutant fish were maintained as previously described^47,53,62,66 44^. *Cyc*^*+/−*^;*oep*^*−/−*^ and *sqt*^*+/−*^;*oep*^*−/−*^ double mutants were generated by incrossing *cyc*^*+/−*^;*oep*^*+/−*^ or *sqt*^*+/−*^;*oep*^*+/−*^ respectively and rescuing them with an injection of 55pg *oep* mRNA at the 1-cell stage. *Smad2*^−/−^ germline carrier fish were obtained by germline transplantation, using *Smad2*^+/−^ incross progeny as germ cell donors^67^. Germline carrier embryos were obtained by either incrossing EK fish or crossing *dmrt1*^E3ins*−/−*^ female fish to *dmrt1*^E3ins*−/+*^ male fish. The *dmrt1*^E3ins*−/−*^ and *dmrt1*^E3ins*−/+*^ fish were gifts from Kaitlyn A. Webster/Kellee R. Siegfried and were used with the intent of biasing germline carriers to female adult fates^68^.

For experiments shown in the text, mutant embryos were derived as follows: *MZoep* embryos were obtained by crossing *oep*^−/−^ adults; *Zoep* embryos were obtained by crossing *oep*^*+/−*^ females with *oep*^*−/−*^ males (see genotyping below); *MZsmad2* embryos were obtained by crossing *smad2*^*−/−*^ germline carrier adults; *Mvg1* embryos were obtained by crossing *vg1*^*−/−*^ females with TLAB males; *lft1*^*−/−*^;*lft2*^*−/−*^ embryos were obtained by crossing *lft1*^*−/−*^;*lft2*^*−/−*^ adults; *sqt*^*+/+*^;*MZoep*, *sqt*^*+/−*^;*MZoep*, and *sqt*^*−/−*^;*MZoep* embryos were obtained by crossing *sqt*^*+/−*^;*oep*^*−/−*^ adults; *cyc*^*+/+*^;*MZoep*, *cyc*^*+/−*^;*MZoep*, and *cyc*^*−/−*^;*MZoep* embryos were obtained by crossing *cyc*^*+/−*^;*oep*^*−/−*^ adults.

### mRNA synthesis and microinjection

pCS2+ vectors containing the CDS of either *SV40NLS-sfgfp* or *oep* were linearized with NotI and subsequently purified with the E.Z.N.A. Cycle Pure (Omega) kit. Purified templates were transcribed using the mMESSAGE mMACHINE SP6 (Invitrogen/Thermo Fisher Scientific) kit, and the resulting *gfp* and *oep* capped mRNAs were purified with the E.Z.N.A. Total RNA Kit I (Omega). Capped mRNA concentrations were evaluated via NanoDrop (Thermo Fisher Scientific) spectrophotometry. Kits were used per manufacturer’s respective protocols.

### Sensor cell transplant experiments

M*vg1* sensor donors were injected with either 1nl of 55pg/nl *gfp* mRNA or 1nl of 55pg/nl *gfp* mRNA+110pg/nl *oep* mRNA (Fig. 3B) at the 1-cell stage. *MZsmad2+oep* hosts (Fig. 3D) were injected with 1nl of 55pg/nl *oep* mRNA at the 1-cell stage. Prior to injection, both donor and host embryos were enzymatically dechorionated using 1mg/ml Pronase (Millipore Sigma). After injection, embryos were raised at 28.5ºC in 1% agarose-coated plastic dishes in embryo medium. At high stage, donor and host embryos were placed in 1X Danieau’s buffer, and ~5-10 blastomeres were transplanted from the animal pole of donor embryos to the margin of host embryos, unless specified otherwise. After transplantation, host embryos were returned to embryo medium and raised to 50% epiboly at 28.5ºC before fixation.

### α-pSmad2 Immunostaining

The protocol was modified from^47^. Briefly, embryos were fixed in 4% formaldehyde overnight at 4ºC in 1x PBSTw (1x PBS + 0.1% (v/v) Tween 20), washed in 1x PBSTw, dehydrated in a MeOH/PBST series (25%, 50%, 75%, and 100% MeOH), and stored at −20°C until staining. Embryos were rehydrated in a MeOH/PBSTr (1x PBS + 1% (v/v) Triton X-100) series (75%, 50%, and 25% MeOH), washed 3x in PBSTr, and manually de-yolked. Embryos were then incubated for 2 hours at room temperature (RT) in antibody binding buffer (PBSTr +1% (v/v) DMSO) before overnight incubation with 1:1000 α-pSmad2 antibody (Cell Signaling Technology #18338) and, when required, 1:1000 α-GFP antibody (Aves Labs AB_2307313) in antibody binding buffer at 4ºC. After 1º antibody incubation, embryos were washed 6X with PBSTr before a 30 min RT incubation in antibody binding buffer. Embryos were then incubated in 1:2000 goat α-rabbit Alexa 647 conjugate (ThermoFisher A-21245) and, when required, 1:2000 goat α-chicken Alexa 488 conjugate (ThermoFisher A-11039) in antibody binding buffer. Embryos were then washed 6X with PBSTr and 1X PBSTw respectively before a 30 min RT incubation with DAPI. Embryos were washed 3X in PBSTr before dehydration in a MeOH/PBSTw series (50% and 100% MeOH). Embryos were stored at −20ºC in MeOH until imaging.

### Embryo clearing and imaging

Embryos were first cleared in 2:1 benzyl benzoate:benzyl alcohol (BBBA)^69^. After clearing, embryos were mounted in BBBA in individual wells of a 15-well multitest slide (MP Biomedicals). Mounting was performed under a Zeiss Stemi 2000 stereoscope fitted with a Nightsea adaptor system with UV filters and light head to enable embryo visualization. Embryos were then cracked with forceps before placement of a #1.5 coverslip, approximately flattening the embryos. The coverslip was secured with adhesive tape before imaging on a Zeiss LSM-700 inverted confocal microscope.

### smFISH probe synthesis

Single molecule fluorescent *in situ* hybridization (smFISH) probes against the coding sequences of *cyclops* and *squint* were drafted using the Stellaris Probe Designer, with oligo length 18-22bp and minimum spacing length 2 nucleotides. Probes were then checked for cross-reactivity between orthologs (probes with <4 mismatches were discarded) and ordered with 3’ C7 amino group modifications (IDT). 39 probes against *cyclops* and 44 against *squint* were purchased. Probe libraries for each gene were pooled together, dehydrated in a Speedvac, and resuspended in water at a concentration of 1mM. Probes were then coupled to Atto-647N NHS ester (Millipore Sigma #18373) per supplier protocol and purified with the Zymo Oligo Clean and Concentrator kit. Probe concentration was then determined using NanoDrop (Thermo Fisher Scientific) spectrophotometry.

### smFISH staining and imaging

The smFISH staining protocol is modified from previous reports^70,71^. Briefly, embryos were fixed in 4% formaldehyde overnight at 4ºC in 1x PBSTw (1x PBS + 0.1% (v/v) Tween 20), washed in 1x PBSTw, dehydrated in a MeOH/PBST series (50% and 100% MeOH), and stored at −20°C until staining. Embryos were rehydrated in a MeOH/PBSTw (50% and 100% PBSTw) series before manual deyolking. Embryos were then incubated in pre-hybridization buffer (preHB) (10% formamide, 2x SSC, 0.1% (v/v) TritonX-100), 0.02% (w/v) BSA, and 2 mM ribonucleoside-vanadyl complex (NEB) for 30 minutes at 30ºC before overnight incubation with 10nM probes in hybridization buffer (10% (w/v) dextran sulfate (MW 500,000) in preHB) at 30ºC in the dark. After staining, embryos were washed 2 x 30 minutes at 30ºC in hybridization wash solution (10% (v/v) formamide, 2x SSC, 0.1% (v/v) Triton X-100) before a brief wash in 2x SSC + 0.1% (v/v) Tween-20. Finally, embryos were incubated for 20 minutes at 30ºC in 0.2X SSC before a 15 minute incubation in DAPI and 2X 2x SSC+0.01%Tween washes.

For membrane staining, 1:100 α-eCdh1 antibody (BD Biosciences #610181) was added overnight with the probes in hybridization buffer. After the 20 minute 0.2X SSC wash, 1:750 Goat α -mouse IgG (H+L)-Alexa 488 (ThermoFisher A32723) in PBSTw was added, and embryos were incubated for 2 hours at RT in the dark. Embryos were washed 6X with PBSTw before a 15 minutes DAPI incubation and 2X 2x SSC+0.01%Tween washes.

For mounting, embryos were kept in 2X SSC, cut from the margin to the animal pole with a scalpel, and mounted in 2X SSC on a standard glass slide between two double-sided adhesive tape bridges (3M Scotch). A #1.5 coverslip then approximately flattens the embryo and is secured in place by the adhesive tape. Mounted embryos were then imaged on a Zeiss LSM-880 inverted confocal using the Airyscan detector.

### Image segmentation

Staining intensities for individual nuclei were compiled for Figs. 1–3. Nuclei were segmented from DAPI channel images using a custom pipeline implemented in MATLAB as described previously^47^. Before segmentation, each image stack was manually inspected to identify acceptable z-bounds. Lower bounds were chosen to exclude internal YSL nuclei from the segmentation. Briefly, for each slice, out-of-plane background signal was approximated by blurring adjacent Z-slices with a Gaussian smoothing kernel and subtracted. Nuclei boundaries were identified using an adaptive thresholding routine (http://homepages.inf.ed.ac.uk/rbf/HIPR2/adpthrsh.htm). Spurious objects were discarded by morphological filtering (area threshold followed by image opening with a disc-shaped structuring element).

Three-dimensional objects were compiled from the two-dimensional segmentation results with a simple centroid-matching scheme. A disc of diameter 5 pixels was defined centered at the centroid of each two-dimensional object, and three-dimensional objects were identified by object labeling with a 6-connected neighborhood. Intuitively, this procedure matches objects whose centroids are separated by <10 pixels (i.e twice the disc diameter used prior to object matching). Objects that fail to span at least 2 Z-slices were discarded. Fluorescence intensities in the DAPI, GFP and pSmad2 channels were compiled as average pixel intensities within the three-dimensional segmentation boundaries.

### Genotyping of Z*oep, cyc;oep* and *sqt;oep* mutant embryos

Crosses leading to homozygous Z*oep*, *cyc;oep* and *sqt;oep* mutant embryos were generated from non-homozygous parents. Specifically, Z*oep* embryos were generated by crossing an *oep*^*−/−*^ male against a oep^+/−^ female; c*yc;oep* embryos were generated from a cross between *cyc*^*+/−*^;*oep*^*−/−*^ parents; *sqt;oep* embryos were generated from a cross between *sqt*^*+/−*^;*oep*^*−/−*^ parents. To identify the genotype of embryos used for imaging, each embryo was manually cut into halves (i.e. through the animal pole) with a clean scalpel after pSmad2 immunostaining. One half of the embryo was dehydrated for clearing and imaging (as described in the α-pSmad2 immunostaining methods section), and the other was used for genomic DNA preparation and genotyping. Genotyping was carried out for each mutation as summarized above. For Z*oep* staining, genotyping was carried out as described for 30% epiboly and 50% epiboly stages; this revealed that Z*oep* embryos could be clearly identified by average staining intensity. Shield-stage Z*oep* embryos were identified by staining intensity.

### Sensor cell identification and gradient quantification

All gradient quantifications in Figs. 1–3 plot nuclear staining intensity as a function of distance from the embryonic margin. Because the margin boundary is curved in our flat mounts, these distances are not a simple function of position within the image. A semi-automated routine was therefore implemented in MATLAB to calculate the distance from the margin for each nucleus. In brief, the YSL-embryo boundary was manually identified and drawn using maximum intensity projections of the DAPI channel. This boundary was then converted into a binary mask and a distance transform was applied. After the distance transform, every pixel in the image adopts a value equal to its distance to the closest non-zero pixel (i.e. the margin contour); the distance from the margin for each nucleus was defined as the pixel intensity of the distance transform image at the corresponding centroid position.

In order to quantify the gradients in Nodal-insensitive host embryos, sensor cells had to be specifically identified. A classification scheme based on nuclear GFP intensity was therefore devised. Because there was some background α-GFP staining, even in cells that did not receive *gfp* mRNA, the approximate baseline GFP intensity was identified by taking a sliding window median of GFP staining intensity as a function of nuclear distance from the margin. GFP^+^ cells were identified as having nuclei brighter than 3-fold above the local baseline, and GFP^−^ cells were identified as having staining intensity at or below the local baseline. These thresholds are stringent and resulted in some false-negative nuclear classifications (e.g. likely GFP^+^ nuclei that failed to be classified as such). However, they do ensure that the nuclei plotted in the main text represent *only* clear GFP^+^ or GFP^−^ populations. This analysis was also performed using less stringent thresholds and manual correction of results, which generated comparable conclusions to the results presented in the paper.

After calculation of GFP staining status and distance from the margin for each nucleus, average gradients were compiled. To facilitate comparison between replicate embryos, the pSmad2 staining intensities were normalized to the baseline intensity (i.e. average nuclear intensity of all nuclei falling between 150 and 200 µm) from the margin. After this normalization, data from each embryo were pooled, and average gradients were compiled with a sliding window average (solid curves in quantified gradients in Figs. 1–3) with a window size of 20 µm. Due to sparse sampling of the gradients by sensor cells, some statistical fluctuations in average gradient shape are evident (e.g. the ‘hump’ in Fig. 2C).

### Kymograph preparation in Figs. 5D and E

In the experimental section of Fig. 5, kymographs were presented that average the behavior of replicate embryos (bars to the right of representative images in Figs. 5D and E). To prepare these kymographs, the distance from the margin for each pixel in the maximum intensity projection α-pSmad2 image was calculated as described in the above section. Pixels were then binned by distance from the margin and averaged across embryos to generate the plots in Fig. 5. Each vertical bar in the plot was drawn for all of the data from a given stage (from left to right: dome, 50% epiboly and shield). Color scalings were selected for visibility and are not equivalent between the wild-type and Z*oep* datasets.

### Gradient simulations

Sensor cell assay simulations were implemented in MATLAB using the PDE toolbox. Simulations were carried out on a two-dimensional rectangular slab (100 x 300 µm) with no-flux boundary conditions. The Nodal source was simulated as a thin strip of tissue (the first 5µm) that produced Nodal at a constant rate. Sensor cells were simulated as small circular domains with permeable boundaries (6 µm diameter) in which parameters (e.g. presence or absence of free receptors) could be set independently of the rest of the tissue. Simulations were run ~2.5 hours of simulation time in an effort to mimic the normal duration of Nodal gradient spread in zebrafish embryos. Simulations are described in detail in the SI (*Reproduction of sensor cell assay with gradient simulations*). Plots in Fig. 4 depict the entire tissue domain at the conclusion of the simulations.

Simulations incorporating receptor production and replacement were implemented in MATLAB using pdepe. Simulations were carried out on a one-dimensional tissue (300 µm long) with no-flux boundary conditions. The Nodal source was simulated as a thin strip of tissue (the first 5 µm) that produced Nodal at a constant rate. Simulations were run for ~2.5 hours of simulation time in an effort to mimic the normal duration of Nodal gradient spread in zebrafish embryos. Simulations are described in detail in the SI (*Gradient simulations accounting for receptor production and consumption*). Plots in Figs. 5B and C are kymographs summarizing the state of the system at regularly sampled times. Each column of kymograph shows the concentration of a given component at each position in the system (‘YSL’ at the bottom), and adjacent columns are separated by 20 s of simulation time. Kymographs begin plotting data at t=0 to capture the transients associated with gradient formation. Pixel scalings (i.e concentration scales) are not identical between Figs. 5b and 5c; scalings were chosen to maximize data visibility. Due to the absence of receptor replacement, concentrations of free receptor and receptor-ligand complexes are markedly lower in Fig. 5c (in accordance with experimental data in Z*oep* mutants, see Fig. 5F).

