## Supplemental Text for "The pattern of Nodal morphogen signaling is shaped by co-receptor expression"

**SI Text**

**Computational Models of Nodal Gradient Formation**

Reproduction of sensor cell assay with gradient simulations

Simulations presented in Fig. 4 were implemented using the MATLAB PDE toolbox. The host embryo was represented as a two-dimensional rectangular slab (100 x 300 µm). 'Sensor cells' were simulated as circular domains of 6 µm diameter—with independently set simulation parameters— scattered throughout the rectangular domain. Equations governing the model are specified as below:

$$\frac{\partial N(x,t)}{\partial t} = \mathrm{D}_N \nabla^2 N(x,t) - k_1 N(x,t) R + k_{-1} C(x,t)$$

$$\frac{\partial C(x,t)}{\partial t} = k_1 N(x,t) R - k_{-1} C(x,t) - k_2 C(x,t)$$

$$\frac{\partial S(x,t)}{\partial t} = k_s C(x,t) - k_{-s} S(x,t)$$

Where $N(x,t)$, $C(x,t)$, and $S(x,t)$ refer to the concentration of free Nodal, Nodal-Receptor complex, and pSmad2 at position $x$ at time $t$, respectively. Parameter values are summarized in the table below. All boundaries are specified as no-flux, and $N(x,t=0)=0$ and $S(x,t=0)=0$ were assumed for initial conditions. The Nodal source was simulated by specifying a constant Nodal production rate ($\lambda_N$) for points lying in the region $0 \leq x \leq 5$. For simplicity, we assume the receptor concentration, $R$, to be constant at each position throughout the simulation. Simulations were run for ~2.5 hours of simulation time to mimic the normal duration of Nodal spread in zebrafish embryos.

Remarks:

1. These simulations instantiate a simple model of morphogen gradient formation—constant synthesis at a localized source coupled with linear degradation—that has been discussed at length elsewhere[1-4]. The steady state gradient for this is a single exponential with length scale set by the diffusion constant and effective Nodal degradation rate (i.e. through complexing with the receptor). Small deviations from this expectation arise due to the finite size of the Nodal source domain.
2. For the plots in the main text, we track signaling through $S$, the concentration of phosphorylated Smad2. We assume the phosphorylation rate to be first-order with respect to the ligand-receptor complex—effectively that unphosphorylated Smad2 is neither depletable nor present in high enough concentrations to reveal saturation of receptor complex kinase activity—and the dephosphorylation rate to be first order with respect to $S$. Intuitively, this means that $S$ reflects receptor occupancy over a time window of $\sim 1/k_{-s}$. Signaling could also be tracked as the concentration of occupied receptor and yields qualitatively similar results.
3. For simulations of sensor cells transplanted into MZ*oep* hosts (*cf* Figs. 2B, 4C), the receptor concentration ($R$) was set to zero outside of the sensor cell boundaries. Within these boundaries, $R$ was kept at the 'wild-type' level (i.e at the same levels as in the simulations from Fig. 4b). We note that the increased signaling intensity in this background reflects the fact that binding with receptor is the only available degradation pathway for the ligand; when $R = 0$, available ligand concentrations are substantially higher throughout the "embryo".
4. For simulations of sensor cells transplanted into *oep*-overexpressing hosts (i.e. MZ*smad2* + *oep* mRNA, *cf* Figs. 3d, 4d) , $R$ was increased by a factor of 2 throughout the host regions, and $k_s$ was

set to zero to mimic the absence of Smad2. Simulation parameters within the sensor cell regions retained their 'wild-type' values.

Gradient simulations accounting for receptor production and consumption

The model presented in Fig. 5 explicitly accounts for production and consumption of receptor components. For the figure panels, the model was simulated on a 1-dimensional domain of length 300 μm. Model equations were specified as follows:

$$\frac{\partial N(x,t)}{\partial t} = \mathrm{D}_N \nabla^2 N(x,t) - k_1 N(x,t)R(x,t) + k_{-1}C(x,t)$$

$$\frac{\partial R(x,t)}{\partial t} = k_3 - k_1 N(x,t)R(x,t) + k_{-1}C(x,t) - k_{-R}R(x,t)$$

$$\frac{\partial C(x,t)}{\partial t} = k_1 N(x,t)R - k_{-1}C(x,t) - k_2 C(x,t)$$

Here, $N(x,t)$, $R(x,t)$, and $C(x,t)$ refer to the concentration of free Nodal, free receptor and receptor-Nodal complexes, respectively, at position $x$ and time $t$. No-flux boundary conditions were assumed at both ends of the domain, and Nodal was produced at a constant rate $\lambda_N$ within a source domain covering $0 \leq x \leq 5$. $N(x,t=0)$ and $C(x,t=0)$ were initiated at zero throughout the field, and $R(x,t=0)$ was set to $\lambda_R/k_{-R}$ for all positions. Simulations were implemented in MATLAB using pdepe.

Remarks:

1. In this model, receptor synthesis is treated as constitutive throughout the field, and degradation occurs through ligand-dependent and ligand-independent mechanisms. The ligand-dependent pathway occurs through degradation of receptor-ligand complexes (schematically represented as endocytosis in Figs. 4 and 5, rate $k_2C(x,t)$). The ligand-independent pathway is assumed to be first-order with respect to free receptor (rate $k_{-R}R(x,t)$).
2. Direct degradation of the receptor—once we have made the step of assuming constitutive production– is required to achieve steady state concentrations of free receptor outside the domain of ligand diffusion. In the absence of this degradation route, receptor levels increase without bound far from the source as the synthesis term is not coupled to receptor levels. We note that this constitutive degradation explains the gradual decrease of free receptor levels far from the source in the Fig. 4c; without replacement, ligand-independent degradation would eventually remove all of the receptor in the system. We include this degradation mechanism in the '*Zoep*' simulations for parity, however the signaling wave occurs even with $k_{-R}$ set to 0.
3. Ligand-dependent removal of receptor is the key requirement for appearance of the signaling wave in *Zoep* mutants. While we regard ligand-dependent endocytosis of receptor complexes as biologically plausible, our model is agnostic to the actual mechanism. Other mechanisms can be imagined; for example, irreversible inactivation of receptor-ligand complexes that remain on the cell surface could also support formation of a wave provided that the ligand does not dissociate. Indeed, we suspect that any mechanism through which ligand binding renders receptors incapable further ligand capture or signaling would support wave formation.
4. For Figs. 5B and C, we summarize the simulation results for each component with kymographs. Each column of these images shows the state of the 1-dimensional system—with source at the bottom and 'animal pole' at the top—at each point. Simulation time proceeds from left to right, and each plot represents two hours of simulation time. We note that, with biologically reasonable

parameters, we observe formation of a stable gradient with receptor replacement and clear propagation of the wave without receptor replacement.

**Model Parameters**

| **Parameter** | **Description** | **Value** | **Intuition for value** | **Reference** |
|---|---|---|---|---|
| $D_N$ | Free Nodal diffusion rate | 30 μm2/sec | MSD of ~650 μm after 2 hours | Muller et al[5] |
| $k_1$ | Nodal-Receptor association rate | 10 μM/sec | Mean time to capture of ~2 sec at initial Oep concentration | DeCrescenzo et al[6] |
| $k_{-1}$ | Nodal-Receptor dissociation rate | 6.25e-4/sec | Mean time to dissociation of 25 minutes | DeCrescenzo et al[6] |
| $k_2$ | Nodal-Receptor complex internalization rate | 1.7e-3/sec | Mean time to internalization of 10 minutes | - |
| $k_3$ | Receptor production rate | 1.6e-5 μM/sec | Approximately 0.7 molecules/sec produced by a cell of 10 μm diameter | - |
| $k_{-R}$ | Receptor degradation rate (ligand independent) | 2.7e-4/sec | Average lifetime of 1 hour | - |
| N(x,t=0) | Free ligand initial condition | 0 μM for all x | Embryo starts out with no Nodal. | - |
| R(x,t=0) | Free receptor initial condition | 0.06 μM for all x | Approximately 2900 molecules for a cell of 10 μm diameter | Dyson and Gurdon[7] |
| C(x,t=0) | Nodal-Receptor complex initial condition | 0 μM for all x | Embryo starts out with no receptor-ligand complexes. | - |

**Primer Table:**

| **Name** | **Sequence** | **Purpose** |
|---|---|---|
| NL-102 | GGGACGGGAGATTCATAGCA | *cyc* forward primer |
| NL-103 | ACACATATACCGTAGTACCTGC | *cyc* reverser primer |
| NL-96 | gccagctgctcgcattttattcc | *sqt* WT primer |
| NL-95 | gagctttatttcaataactgcgtg | sqt common primer |
| NL-94 | atataaaatcagtacaaccgcccg | *sqt* mutant primer |
| NL-89 | GCAAAGAAGATATGCACTCAATCATTTATTAAATGG | *smad2* forward primer |
| NL-91 | GGGCTCTGAACAAAGATGGCG | *smad2* reverse primer |
| AC102 | AGGCCCTCGAGATAAATAACA | *oep* forward primer |
| AC103 | ACAGCAAACATCAAGAACCTG | *oep* reverse primer |
| AC149 | CTTGCAGATGTGGATTTCTGGCC | *vg1* forward primer |
| AC150 | CATGATGCGATGGTTTGGGTCG | *vg1* reverse primer |
| NL-36 | catgtatcaccttccctctgatgtc | *lft1* forward primer |
| NL-37 | gcattagcctatatgttaacttgcac | *lft1* reverse primer |
| NL-99 | CATTTTGACCACAGCGAT | *lft2* WT allele forward primer |
| NL-100 | GTTCATTTTGACCACTCAC | *lft2* mutant allele forward primer |
| NL-101 | gaattgtgcataagtaacccacctg | *lft2* common reverse primer |
