## Supplementary Figures for "The pattern of Nodal morphogen signaling is shaped by co-receptor expression"

**FIGURE S1**

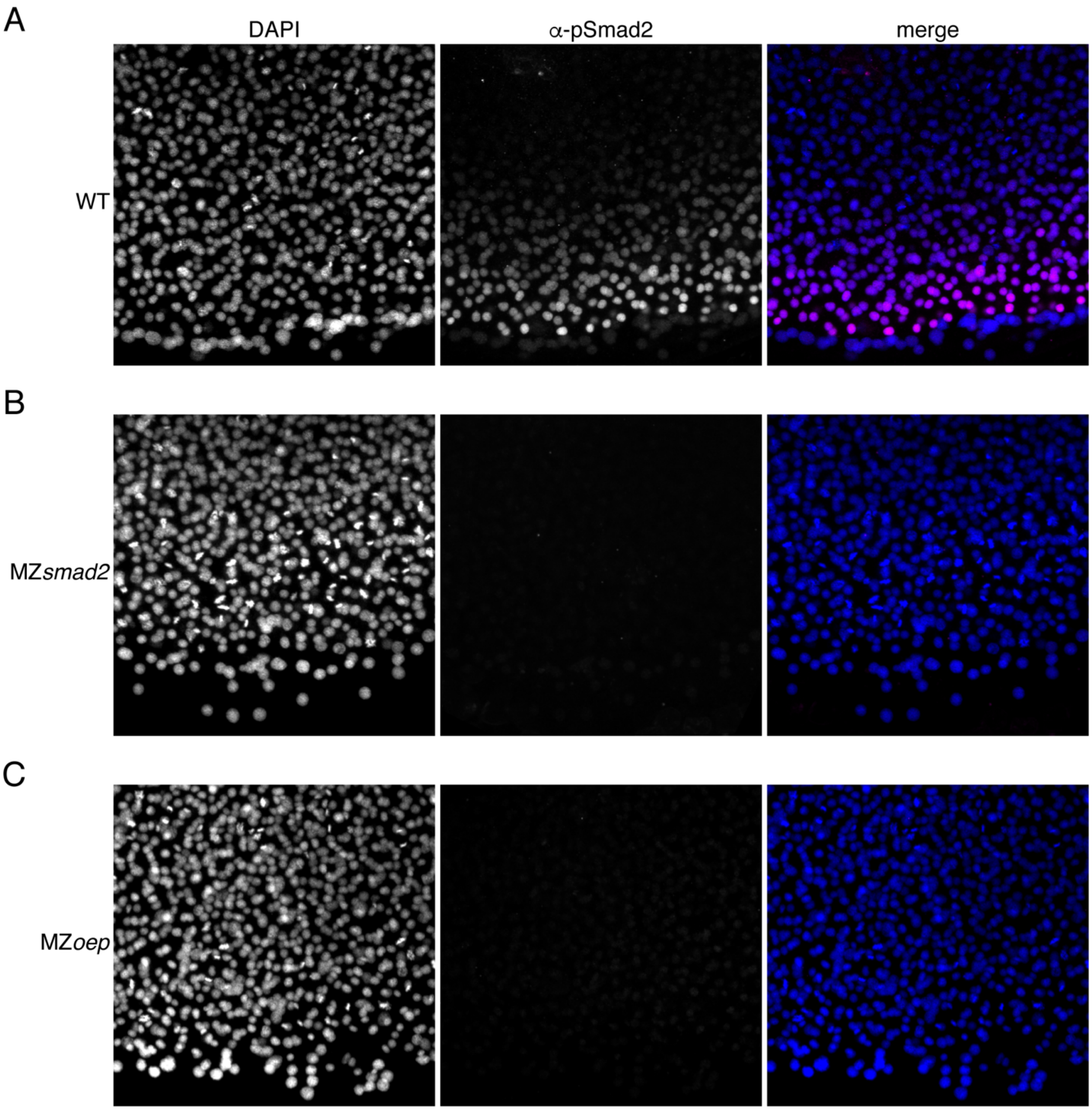


**<u>Fig. S1. MZ*smad2* an MZ*oep* mutants lack pSmad2.</u>** This experiment summarizes control experiments that verify that our α-pSmad2 staining protocol detects Nodal signaling activity in wild-type embryos but not MZ*smad2* and MZ*oep* mutants. **A)** Flat-mount image of wild-type 50% epiboly embryo stained with DAPI and α-pSmad2 antibody. Images are maximum intensity projections from a representative embryo. **B)** Flat-mount image of MZ*smad2* 50% epiboly embryo stained with DAPI and α-pSmad2 antibody. Images are maximum intensity projections from a representative embryo.

**C)** Flat-mount image of MZ*oep* 50% epiboly embryo stained with DAPI and α-pSmad2 antibody. Images are maximum intensity projections from a representative embryo.

**FIGURE S2**

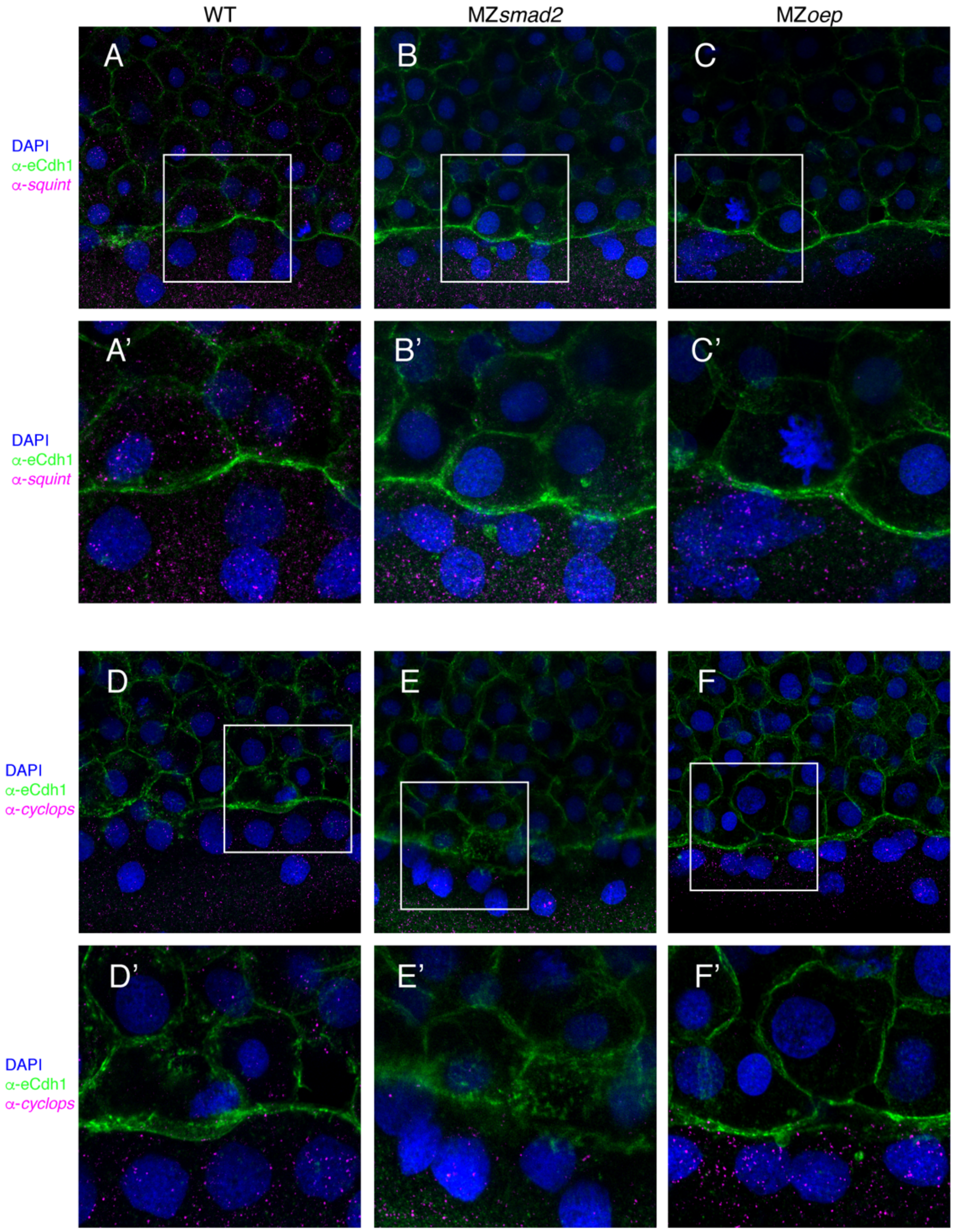


**<u>Fig. S2. MZ*smad2* and MZ*oep* embryos have intact Nodal sources.</u>** To verify that MZ*smad2* and MZ*oep* embryos express Nodals in the YSL, we stained for *cyclops* and *squint* mRNA by smFISH. All depicted embryos were counterstained with DAPI to mark nuclei and α-eCdh1 to mark cell boundaries **A)** Wild-type embryos express *squint* mRNA in the YSL and blastoderm at 50% epiboly. **A')** Enlarged view of area within the white box from panel (A). **B)** MZ*smad2* embryos express *squint* mRNA in the YSL, but not the blastoderm, at 50% epiboly. **B')** Enlarged view of area within the white box from panel (B). **C)** MZoep embryos express *squint* mRNA in the YSL, but not the blastoderm, at 50% epiboly. **C')** Enlarged view of area within the white box from panel (C). **D)** Wild-type embryos express *cyclops* mRNA in the YSL and blastoderm at 50% epiboly. **D')** Enlarged view of area within the white box from panel (D). **E)** MZ*smad2* embryos express *cyclops* mRNA in the YSL, but not the blastoderm, at 50% epiboly. **E')** Enlarged view of area within the white box from panel (E). **F)** MZoep embryos express *cyclops* mRNA in the YSL, but not the blastoderm, at 50% epiboly. **F')** Enlarged view of area within the white box from panel (F).

**FIGURE S3**

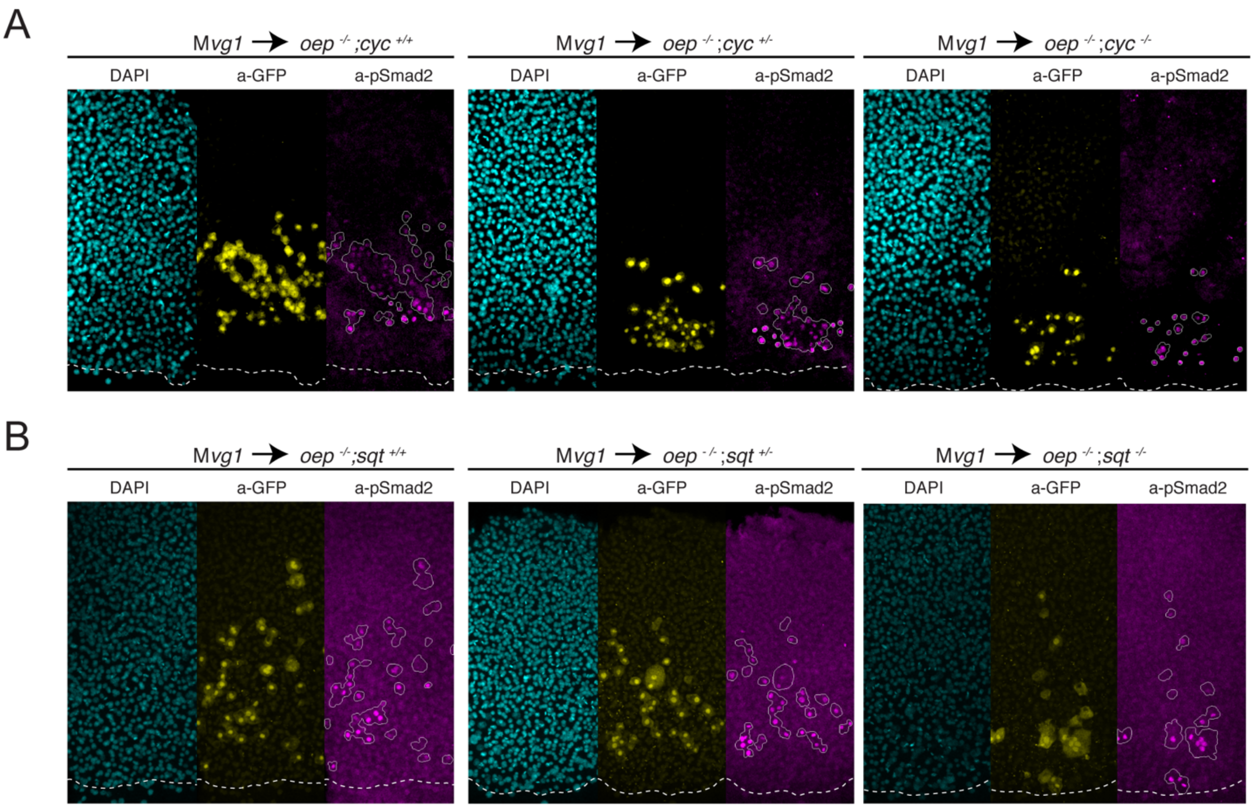


**Fig. S3. Cyclops and Squint signal over a long range in the absence of Oep.** To test whether loss of Oep expands the range of both Cyclops and Squint, we generated MZ*oep;*cyc and MZ*oep;sqt* double mutants and performed sensor cell assays. **A)** Squint signals over a long range in the absence of Oep. Representative sensor cell assays for MZ*oep*;*cyc*$^{+/+}$ (left), MZ*oep*;*cyc*$^{+/-}$ (middle) and MZ*oep*;*cyc*$^{-/-}$ (right) are presented. M*vg1* sensor cells are marked with α-GFP immunostaining (yellow), and sensor cell boundaries are outlined in white in the α-pSmad2 images (magenta). YSL boundaries are marked with a white dashed curve in all images. **B)** Cyclops signals over a long range in the absence of Oep. Representative sensor cell assays for MZ*oep*;*sqt*$^{+/+}$ (left), MZ*oep*;*sqt*$^{+/-}$ (middle) and MZ*oep*;*sqt*$^{-/-}$ (right) are presented. M*vg1* sensor cells are marked with α-GFP immunostaining (yellow), and sensor cell boundaries are outlined in white in the α-pSmad2 images (magenta).
